# Fluorescent polysome profiling reveals stress-mediated regulation of HSPA14-ribosome interactions

**DOI:** 10.1101/860833

**Authors:** Anatoly Meller, David Coombs, Reut Shalgi

## Abstract

Ribosome-associated factors play important roles in regulation of translation in response to various physiological and environmental signals. Here we present fluorescent polysome profiling, a new method that provides simultaneous detection of UV and fluorescence directly from polysome gradients. We demonstrate the capabilities of the method by following the polysome incorporation of different fluorescently tagged ribosomal proteins in human cells. Furthermore, we used fluorescent polysome profiling to examine chaperone-ribosome interactions, and characterized their changes in response to proteotoxic stress conditions. We revealed dynamic regulation of HSPA14-polysome association in response to heat shock, with a marked heat shock-mediated increase in the polysome-association of HSPA14. Our data suggest a model whereby HSPA14 dimerization is increased upon heat shock. We therefore established fluorescent polysome profiling as a streamlined method, that can enhance and facilitate the broad exploration of ribosome-associated factors and their regulation.

## Introduction

Regulation of translation, allowing fast changes in protein levels, is necessary for a timely cellular response to changing environmental or internal conditions (Spriggs, Bushell et al., 2010). Translation regulation can occur globally, through control of various translation factors involved in the different stages of translation. i.e. initiation, elongation or release factors (Dever & Green, 2012, Sonenberg & Hinnebusch, 2009). Translation can also be controlled specifically, at the level of single or groups of mRNAs, or at the level of nascent chains, often through ribosome associated factors (Harvey, Smith et al., 2018, Preissler & Deuerling, 2012). The ribosome itself is also increasingly being implicated in regulation of translation, with specialized ribosomes interacting differentially with specific subsets of mRNAs (Gilbert, 2011, Xue & Barna, 2012).

A commonly used method for assessing global translation levels and changes is polysome profiling, which involves separation of polyribosome complexes by density using sucrose gradients centrifugation and measuring the optical density along the gradient. The resulting curve allows to measure the amounts of RNA bound by mono- and poly-ribosomes, which is used as a proxy for the overall translation levels in the cell.

A large variety of ribosome-interacting proteins have been demonstrated to affect translation either globally, or for specific mRNAs. Examples include the microRNA machinery, various RNA binding proteins, and more (Harvey et al., 2018, Simsek, Tiu et al., 2017), highlighting the importance of studying the specific functions and the dynamic interactions of different types of ribosome-associated factors.

Molecular chaperones have long been known to interact with ribosomes, and aid in the holding of nascent chains and folding of newly synthesized proteins (Pechmann, Willmund et al., 2013, Preissler & Deuerling, 2012). More recently, accumulating evidences suggest that chaperones, being the sensors of the protein homeostasis state in the cell, can play an active role in regulation of translation in response to perturbations in proteostasis (Brandman, Stewart-Ornstein et al., 2012, Kirstein-Miles, Scior et al., 2013, Liu, Han et al., 2013, Shalgi, Hurt et al., 2013). Indeed, we and others have previously shown that HSP70-ribosome association is reduced in response to proteotoxic stress in mouse and human cells (Liu et al., 2013, Shalgi et al., 2013). The ribosome association of the NAC chaperone (Nascent chain Associated Complex) has been shown to be reduced in response to protein aggregation conditions in nematodes (Kirstein-Miles et al., 2013). Therefore, the dynamics and regulation of ribosome-associated factors, including chaperones, is of growing interest.

Nevertheless, the dynamics of the ribosome association of different proteins is often difficult to follow. The most widely used method to assay the ribosome-association of proteins has been fractionation of polysome profiles, followed by protein extraction from the different gradient fractions, and subsequent western blotting. This method is laborious, especially when looking to compare different conditions. More recently, fluorescence monitoring of polysome fractions was performed after fractionation, using fluorescently tagged ribosomal proteins in *E.coli*, to examine ribosome assembly intermediates (Nikolay, Schloemer et al., 2014), improving the efficiency of the traditional method.

Here we present a new technology called fluorescent polysome profiling, which facilitates an easy and streamlined interrogation of polysome-associated factors. This method uses a new flowcell instrument that provides an integrated, simultaneous measurement fluorescence together with UV (260nm, for RNA absorbance), which is combined with a polysome gradient station. We used the instrument to measure UV and fluorescence from polysome profiles of lysates from human HEK293T cells expressing a variety of GFP-tagged proteins, in order to monitor their continuous co-migration with polysomes. This allowed us to obtain high resolution measurements of polysome-association for different proteins, first ribosomal proteins (RPs) and then chaperones, and examine their dynamic association in response to proteotoxic stress conditions.

## Results and Discussion

### Fluorescent polysome profiling – a new method for detection and quantification of polysome-associated proteins

Here we describe Fluorescent polysome profiling - a new method for detection and quantification of the polysome-association of any ribosome associated factor of interest, or the polysomal distribution of ribosome components, performed simultaneously with the generation of the traditional UV-based polysome profiles. The method relies on the Triax™ flowcell, an instrument that provides an integrated, simultaneous measurement of UV (260nm, for RNA absorbance) and fluorescence directly from gradients.

In order to determine that the flowcell is able to reliably detect ribosome-associated fluorescent proteins, we tested its capabilities using fluorescently-tagged RPs. We first cloned a GFP tag to the ribosomal protein RPLP0 (see Methods), and transfected it into human HEK293T cells. GFP-tagged RPLP0 showed a diffused fluorescence pattern in the cytosol (Fig. 1A). Two days after transfection, we performed fluorescent polysome profiling, and examined the fluorescence signal obtained from the polysome gradients, simultaneously with the UV profile, which provides the levels of RNA at each point in the gradient. We note that the ectopic expression of GFP tagged RPLP0 did not affect the level of translation, as judged by completely overlapping polysome profiles of GFP-only and GFP-tagged RPLP0-transfected cells (Fig. S1A). In parallel, we ran fluorescent polysome profiling on non-transfected cells, and on lysis buffer only. Both polysome profiles of GFP-expressing and GFP-tagged RPLP0-expressing cell lysates were very similar to the profile of non-transfected cells, indicating that translation was overall not affected by the transfection procedure, and by the ectopic expression of fluorescently tagged proteins (Fig. S1B). Both the standard polysome lysis buffer (see Methods) and non-transfected cell lysates gave a low fluorescence background along the gradient (Fig. S1C). It is therefore essential to subtract this background from profiles of fluorescently-tagged factors of interest, and thus, in all subsequent experiments, a non-transfected cell lysate sample profile was subtracted from the fluorescence profiles of each GFP-tagged protein that was tested (as illustrated in Fig. S1C,D, see Methods).

**Figure 1:**
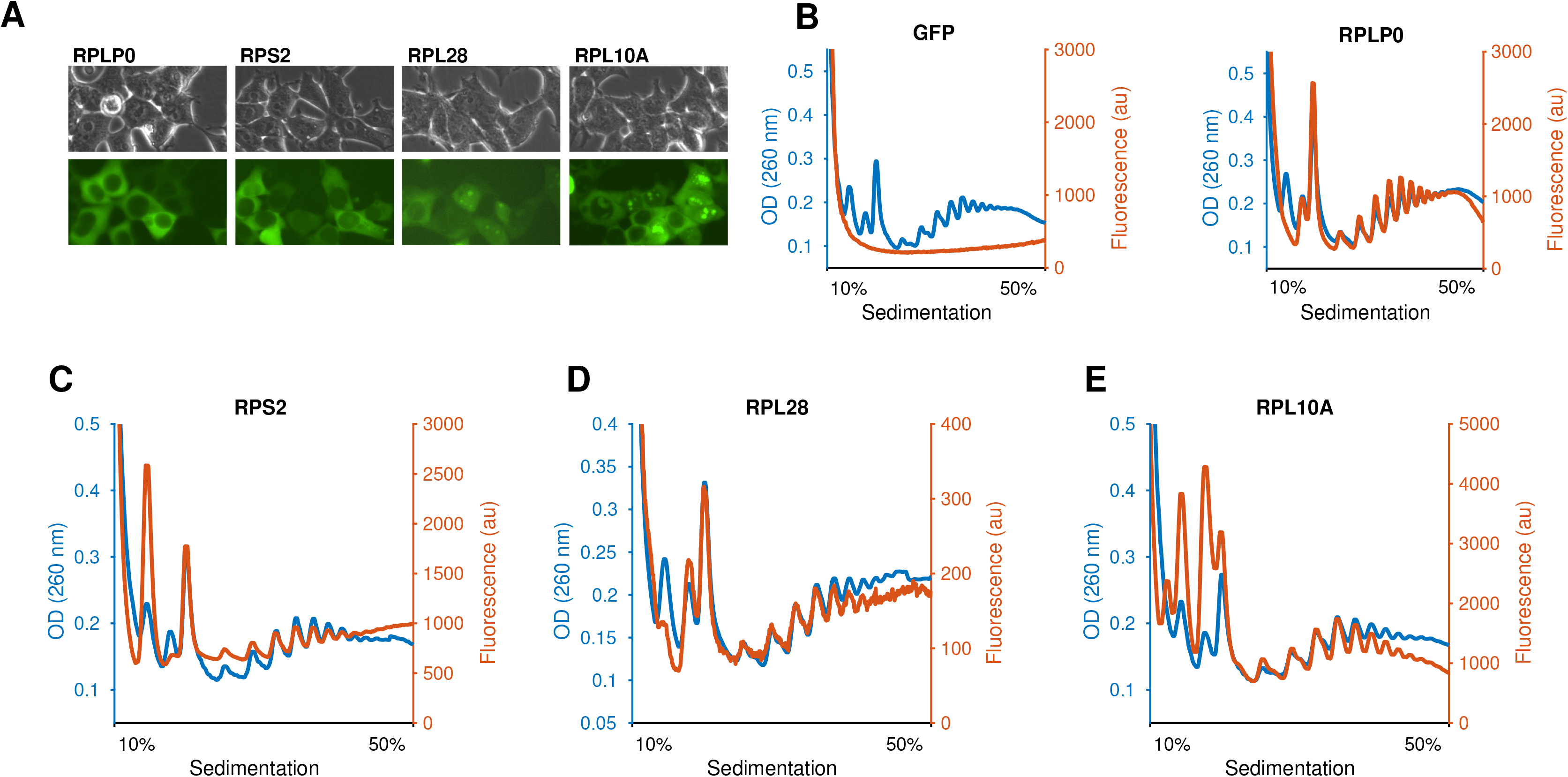
Fluorescent polysome profiling reliably detected GFP-tagged polysome-incorporated ribosomal proteins. (A) Fluorescence and bright field imaging of GFP-tagged ribosomal proteins, expressed in human HEK293T cells, was performed 2 days after transfection with the following exposure times for the GFP channel: RPLP0: 200 ms, RPS2: 200 ms, RPL28: 1000 ms, RPL10A: 400 ms. (B) Fluorescent polysome profiling confirms the specific detection of polysome-incorporated GFP-tagged RPLP0. Free GFP protein remained in the unbound fraction of the polysome curve (orange curve, left panel), while the fluorescence profile of GFP-tagged RPLP0 lysates (orange curve, right panel) closely followed that of the UV, RNA-based, polysome profile (blue curve). (C-E) GFP-tagged ribosomal proteins RPS2 (C), RPL28 (D) and RPL10A (E) all showed ribosome incorporation using fluorescent polysome profiling.

As shown in Fig. 1B, while GFP alone concentrated at the top of the gradient (in the unbound fraction), indicating it did not bind ribosomes, RPLP0 fluorescence profile traced the polysome profile almost perfectly, with the exception of the small subunit (40S) peak (see Fig. S1E,F for an additional biological replicate). Indeed, as RPLP0 is a large subunit ribosomal protein, we do not expect to see it in the 40S peak. This indicated that the fluorescent polysome profiling method reliably detected ribosome-incorporated proteins.

### Fluorescent polysome profiling can be used with different ribosomal proteins

Next, we GFP-tagged additional RPs, and examined their polysome incorporation using fluorescent polysome profiling. We note that all chosen RPs, including RPLP0 used above, are surface accessible (see RPs marked on the ribosome structure in Fig. S2). The different RPs were expressed at different levels (see Methods, Fig. 1A, S1J). RPS2, a small subunit ribosomal protein, gave a strong fluorescence peak at the 40S subunit, as well as at the 80S monosome peak, and significant fluorescent peaks at the various polysomal fractions (Fig. 1C). Importantly, RPS2 showed a negligible peak at the large ribosome subunit (60S) area of the gradient (Fig. 1C, see Fig. S1G for an additional biological replicate). This further confirms the specificity of fluorescent polysome profiling.

GFP-tagged RPL28 was relatively lowly expressed (Fig. S1J), yet the instrument was still able to detect its ribosomal incorporation, demonstrating the sensitivity of the method. The fluorescence profile of GFP-tagged RPL28 largely followed that of the polysome profile, with the exception of the 40S peak, again, as expected from a large subunit RP (Fig. 1D, see Fig. S1H for an additional biological replicate). Finally, RPL10A, which has been previously used for ribosome pulldown assays (Heiman, Schaefer et al., 2008), showed a strong fluorescence profile (Fig. 1E, see Fig. S1I for an additional biological replicate). RPL10A profiles showed several strong peaks at the pre-80S region of the gradient, in addition to the expected 60S subunit peak. Using microscopy imaging, we observed, in addition to the diffused cytoplasmic pattern, strong nuclear punctae consistent with nucleoli localization (Fig. 1A). Thus, it is possible that the pre-80S peaks observed in the fluorescent polysome profiles correspond to different stages in the assembly of ribosome subunits. Future studies will determine the nature of these peaks.

### Investigating ribosome-associated chaperones regulation using fluorescent polysome profiling

We next decided to use fluorescent polysome profiling to examine the ribosome-association of different chaperones. Chaperones are known to associate with ribosomes, and with newly synthesized nascent chains (Pechmann et al., 2013). We focused on the two chaperone systems known to interact with ribosomes, namely the Nascent polypeptide-Associated Complex (NAC) and the Hsp70-mRAC system (Preissler & Deuerling, 2012, Raue, Oellerer et al., 2007, Zhang, Sinning et al., 2017). The NAC chaperone is a heterodimer, composed of NACA and BTF3, that associates with the ribosome and interacts with nascent polypeptides as they emerge from the ribosome exit tunnel (Raue et al., 2007). The mammalian Ribosome Associated Complex (mRAC) co-chaperone system is a ribosome-specific co-chaperone system for the general cytosolic Hsp70s (Hundley, Walter et al., 2005, Jaiswal, Conz et al., 2011, Otto, Conz et al., 2005), while they are recruited to ribosomes and interact with nascent chains (Beckmann, Mizzen et al., 1990, Frydman, Nimmesgern et al., 1994, Nelson, Ziegelhoffer et al., 1992). The mRAC system is comprised of the HSP40 family member MPP11, also known as DNAJC2, a cochaperone that is essential for the stimulation of the ATPase activity and chaperone action of Hsp70, and the HSP70 like protein HSPA14, also known as HSP70L1 (Hundley et al., 2005, Jaiswal et al., 2011, Otto et al., 2005). mRAC associates with the ribosome via MPP11 (Lee, Sharma et al., 2016, Leidig, Bange et al., 2013), and together with Hsp70 interacts with the nascent polypeptide.

We therefore performed fluorescent polysome profiling on cells transfected with either GFP-tagged human BTF3, the beta subunit of the NAC chaperone, or human HSPA14, a subunit of mRAC. Furthermore, as many chaperones are known to play central roles in the regulation of protein homeostasis in response to proteotoxic stresses (Richter, Haslbeck et al., 2010), we wanted to examine potential regulation of the polysome-association of each of these chaperones in response to different perturbations. To that end, we performed fluorescent polysome profiling on transfected cells subjected to each of three proteotoxic stress conditions: heat shock, proteasome inhibition, or ER stress (see Methods).

GFP-tagged BTF3 showed a robust polysome-association under normal growth conditions (Fig. 2A, independent biological replicate presented in Fig. S3A). During stress, BTF3 polysome association changed, in a manner that largely followed the global polysome collapse typical for many stress conditions (Fig. 2B-D, independent biological replicate presented in Fig. S3B-D). Quantification of the changes in BTF3-ribosome association along the gradient indeed showed no significant differences (Fig. S3E). Thus, the NAC chaperone did not display specific regulation of its ribosome association in response to the proteotoxic stress conditions examined here.

**Figure 2:**
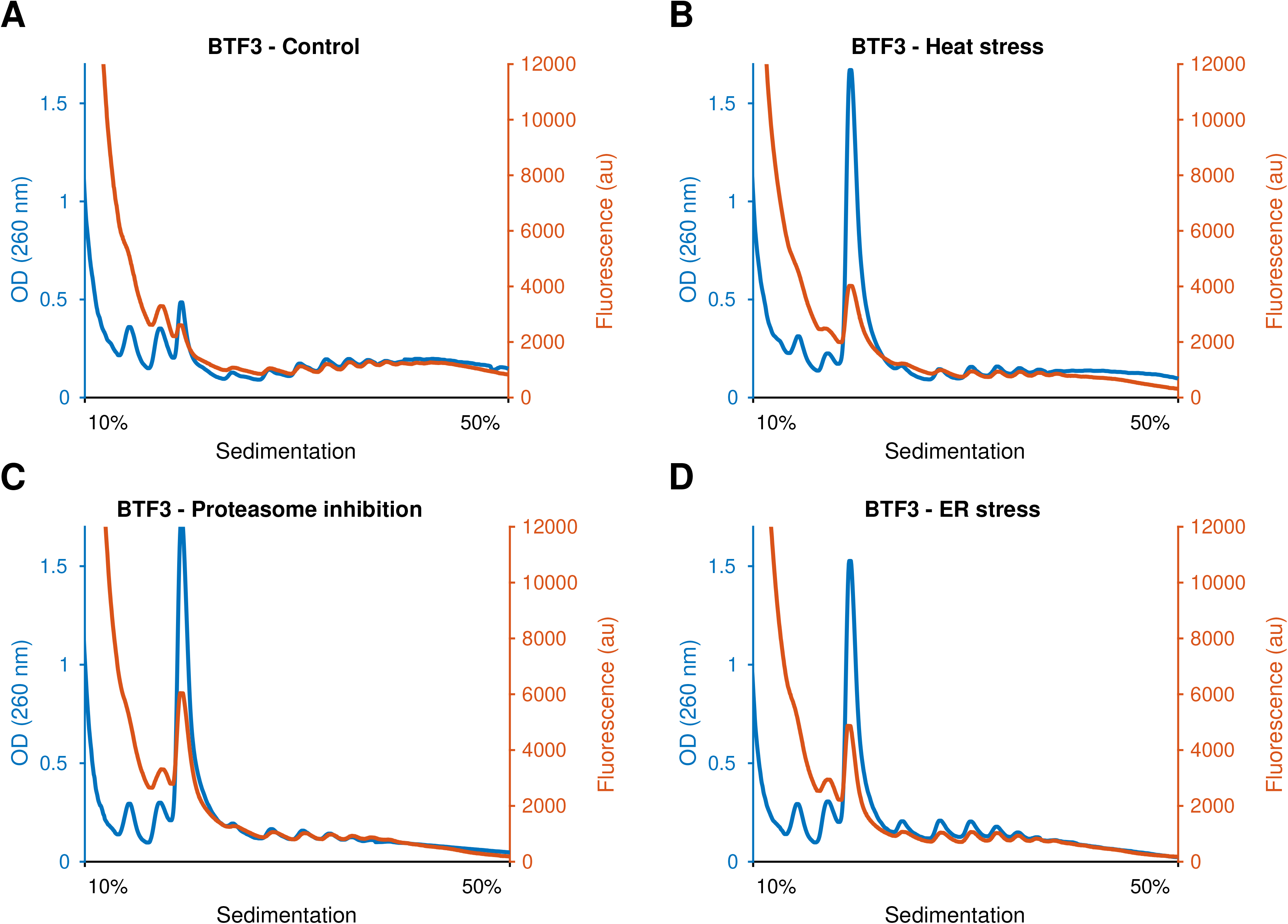
Fluorescent polysome profiling of NAC chaperone-polysome interactions. (A) GFP-tagged BTF3 showed a robust polysome association. (B-D) During stress, BTF3 polysome association (orange curve) largely followed the global polysome collapse typical of translation inhibition, as seen by the UV-based profiles (blue curves), which occur after stress. (B) 2h of heat shock (C), proteasome inhibition (using MG132) (D) and ER stress (using Thapsigargin). Quantification (Fig. S3E) indeed showed no significant differences.

### Fluorescent polysome profiling revealed HSPA14 polysome-association induction following heat shock

Interestingly, examination of the fluorescent polysome profiles of GFP-tagged HSPA14 revealed that its ribosome association was markedly increased following heat shock (Fig. 3). Quantification showed that the association of HSPA14 was highly induced along the gradient (Fig. 3E, see Methods). Moderate increase in HSPA14-ribosome association was also observed following proteasome inhibition (Fig. 3C,E). Previously, we and others showed that Hsp70 ribosome association (specifically, the HSC70 family member) was reduced in heat shock (Shalgi et al., 2013) and other proteotoxic stresses (Liu et al., 2013). As HSPA14 is a subunit of the mRAC, that serves as co-chaperone for the cytosolic Hsp70s when they associate with the ribosome (Pechmann et al., 2013), we initially expected to see ribosomal depletion of HSPA14 during heat stress, similar to that previously shown for Hsp70 (Liu et al., 2013, Shalgi et al., 2013). Nevertheless, our results showed the opposite trend.

**Figure 3:**
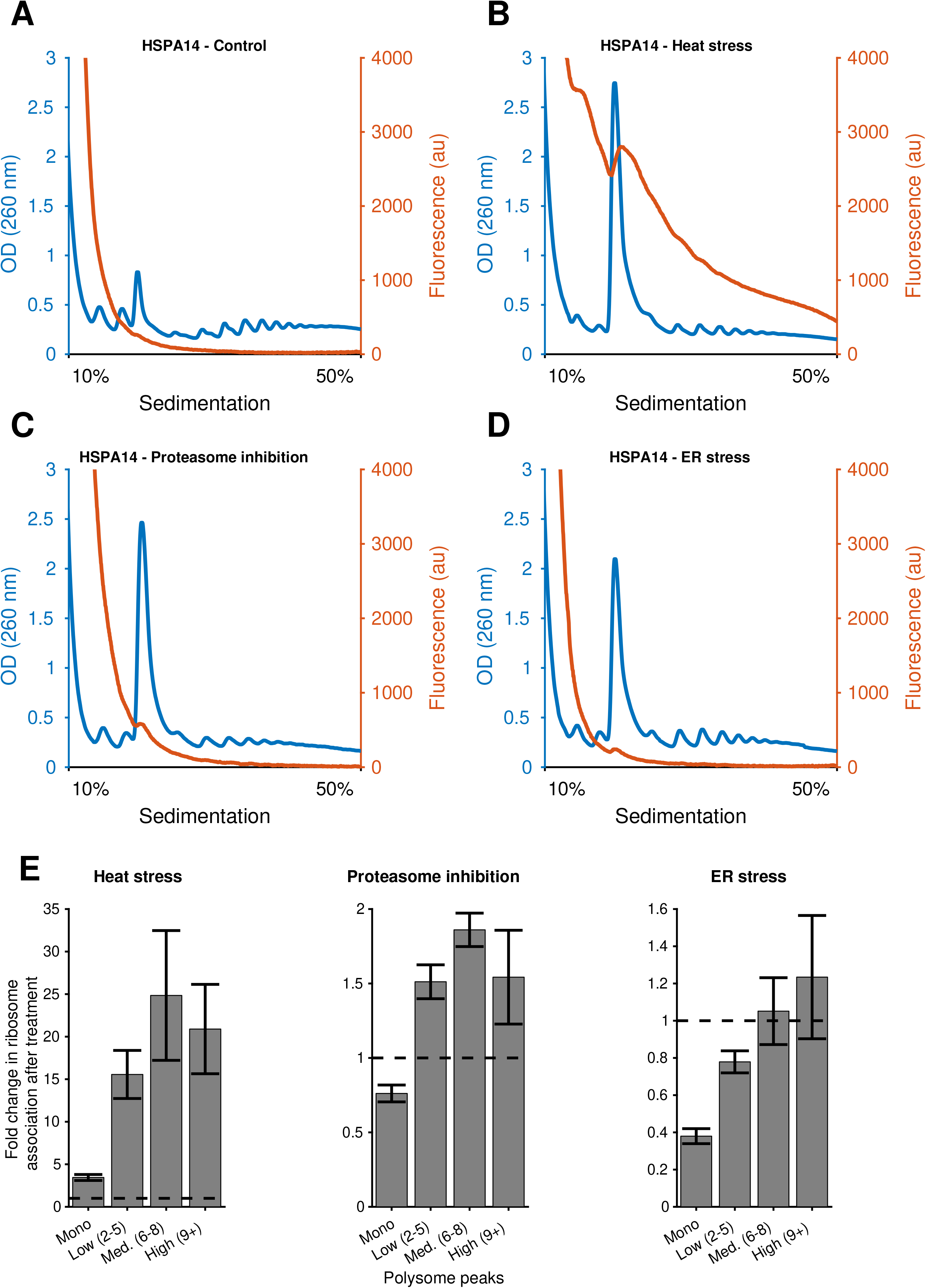
HSPA14 polysome association is induced during Heat stress. (A-D) Examination of an N-terminally GFP-tagged HSPA14 using fluorescent polysome profiling in control and stress conditions (as in Fig. 2). HSPA14 showed a marked increase in polysome association following heat stress (B) compared to control conditions (A). (E) Quantification of the fold change in ribosome-association of HSPA14 showed that the association of HSPA14 was highly induced along the gradient following Heat Stress (see Methods).

To verify that this result was independent of the position of the GFP tag, we generated, in addition to the N-terminal GFP-tagged HSPA14 (Fig. 3), a C-terminal GFP-tagged version of the protein. Here too, we observed the heat shock-mediated induction in the polysome association of HSPA14 (Fig. 4A,B, S4A,B). To further validate this finding, and verify that the heat shock-induced polysome-association of HSPA14 was not affected by the relatively large GFP tag, we transfected cells instead with a FLAG-tagged HSPA14, and assayed HSPA14 polysome-association using the traditional method, namely polysome profiling followed by polysomal fraction collection and western blot (WB). As shown in Fig. 4C (independent biological replicate presented in Fig. S4C), this traditional method indeed demonstrated the marked increase in HSPA14 polysome-association following heat shock, providing further corroboration to the dynamic regulation of HSPA14-polysome association during heat shock.

**Figure 4:**
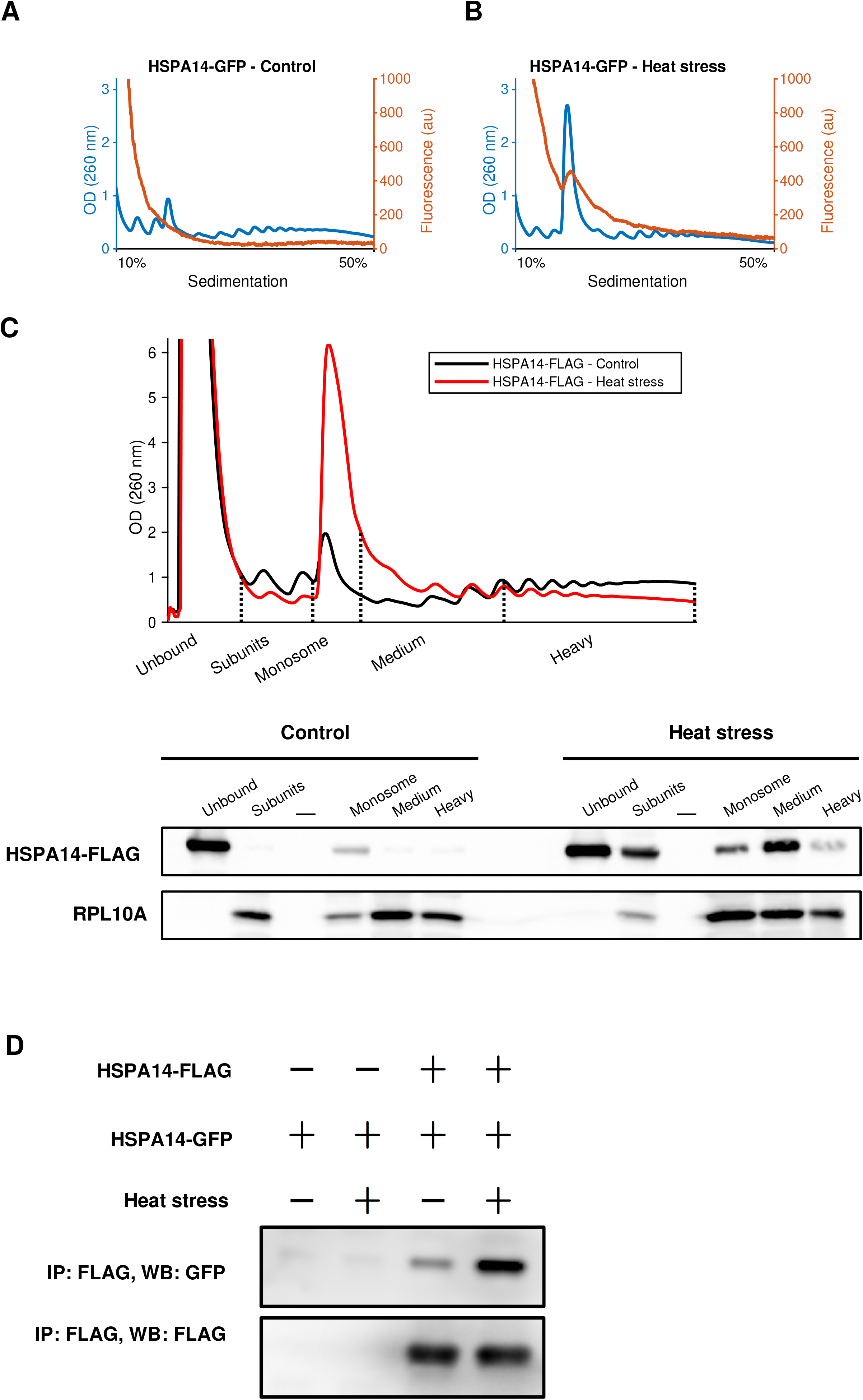
Induced HSPA14 polysome association in heat stress was accompanied by an increased HSPA14 dimerization. (A-B) C-terminal GFP-labeled HSPA14 showed a marked increase in polysome interactions following heat stress (B) compared to control conditions (A), similarly to the N-terminal tagged HSPA14 shown in Fig. 3. See also Fig. S4A,B for more stress conditions and quantifications. (C) Polysome profiles of cells transfected with HSPA14-FLAG show an expected polysome collapse following heat stress (top panel). X-axis dashed lines indicate fractionation of sucrose gradients for protein extraction (top panel, see Methods). Western blot of HSPA14-FLAG protein levels (using anti-FLAG antibody) across the polysome gradient in control and heat stressed cells across the different fractions, RPL10A (endogenous) was used as a ribosomal marker (bottom panel). Unbound fraction showed a similar amount of HSPA14-FLAG in both control and heat stress, and contain no detectible RPL10A, as expected. All other fractions showed a significant increase of HSPA14-FLAG in heat stress, further corroborating our observation obtained using fluorescent polysome profiling (Fig. 3, 4A,B). See Fig. S4C for an independent biological replicate. (D) Coimmunoprecipitation revealed an increased HSPA14-HSPA14 interaction following heat stress. HEK293T cells were co-transfected with FLAG-tagged HSPA14 and GFP-tagged HSPA14 and further subjected to 2h of heat shock. Then immunoprecipitation was performed with anti-FLAG coated beads, and HSPA14-HSPA14 association was inspected using western blot against GFP. Cells not expressing FLAG-tagged HSPA14 have a minimal GFP background, confirming specificity (lanes 1,2). After heat shock, HSPA14-HSPA14 interaction is highly increased compared to control cells (lane 4 compared to 3). See also Fig. S4D-H.

Hsp70 has been previously shown to form dimers. Dimerization of Hsp70 was observed for both *Homo sapiens* and *E.coli* Hsp70 (Morgner, Schmidt et al., 2015, Qi, Sarbeng et al., 2013, Thompson, Bernard et al., 2012), although the functional role of the dimers is still not clear. Since HSPA14 is an Hsp70 homolog, we hypothesized that the observed increase in its association with polysomes might be explained, at least in part, by an increased tendency of HSPA14 to homo-dimerize in heat shock. In order to test this hypothesis, we performed co-immunoprecipitation (co-IP) experiments of co-transfected FLAG-tagged HSPA14 and GFP-tagged HSPA14, in either untreated cells, or cells subjected to heat shock treatment. Interestingly, we observed a significant increase in HSPA14-HSPA14 interactions following heat shock (Fig. 4D, S4D-G, see Methods). This increase in HSPA14-HSPA14 interaction was unchanged when we pre-treated lysates with RNase (Fig. S4H, see Methods), indicating that the observed HSPA14-HSPA14 interactions cannot be the result of indirect association through neighboring ribosomes on the same polysome. Therefore, these results suggest a model in which HSPA14 has an increased tendency to form dimers upon heat shock. Future studies will further illuminate the mechanism and regulation of HSPA14 increased polysome association, and its potential effect on stress-mediated translational control.

## Conclusions

Together, our experiments established fluorescent polysome profiling as a sensitive, high resolution method for the detection and quantification of the polysome-association of ribosome-interacting factors. Using this method, we illuminated novel aspects of the dynamics and regulation of ribosome-associated chaperones, in particular, a novel behavior of ribosome-associated HSPA14 in heat stress conditions. Consequently, fluorescent polysome profiling can be used to explore virtually any ribosome-associated factor that can be fluorescently tagged, and reveal new biological insights into the dynamics and regulation of such factors at the ribosome. This easy and streamlined method can potentially be used to replace the traditional, laborious, method thus far used in the field (i.e. polysome fractionation followed by fraction collection to isolate proteins and their detection using WB). Additionally, it will enable the study of a wide variety of proteins for which a reliable antibody is lacking. Furthermore, a plethora of resources of cells harboring GFP tagged proteins have been developed over the years, from the widely used yeast libraries (Huh, Falvo et al., 2003) to GFP-tagged protein mammalian cell line collections (Harikumar, Edupuganti et al., 2017, Sigal, Milo et al., 2006), and have thus far been extensively used to study protein expression dynamics, localization, and more. Combining these resourced together with fluorescent polysome profiling will potentially facilitate and promote a much broader examination of many additional ribosome-associated factors and their regulation.

Our experiments illustrate the variety of potential applications of fluorescent polysome profiling. These applications can range from the characterization of ribosome composition, ribosome assembly intermediates, ribosome-associated factors, mRNA-associated factors, and more. Utilization of a variant of the flowcell with two different fluorescent detectors will also expand the capabilities of the method, allowing to simultaneously assay two ribosome-associated proteins at the same run.

Fluorescent polysome profiling can also be potentially used to study the polysomal distribution of specific mRNAs, either using fluorescent probes, or through labeling of mRNAs via tethered coat proteins such as the MS2 system (Bertrand, Chartrand et al., 1998). These future applications will require additional calibration steps.

Finally, with the growing interest in specialized ribosomes (Gilbert, 2011, Xue & Barna, 2012), which may have important implications on mRNA-specific translation regulation, fluorescent polysome profiling has the potential to greatly enhance our understanding of the regulation of ribosome composition in different conditions.

## Materials and Methods

### Cell culture and Transfection

HEK293T cells were grown in standard DMEM supplemented with 10% FBS. For transfection, HEK293T cells (5*10^6^ cells per plate) were seeded on 10cm plates and transfected using Lipofectamine 2000 reagent on the same day, according to the manufacturer instruction. As GFP alone was much more highly expressed than GFP-tagged proteins, the amount of transfected DNA for the GFP-only plasmid was titrated down such that the fluorescence peak at the unbound fraction matched that of GFP-tagged RPLP0. Twenty-four hours after transfection, cells were split into two 10cm plates. Cells were harvested 48 hours following the original transfection, when they were at approximately 70% cell confluence. Importantly, we made sure to avoid cell crowding as this inhibits overall cellular translation.

### Proteotoxic stresses

HEK293T cells transfected with GFP-tagged proteins were subjected to the following proteotoxic stresses: Heat shock: 2 hours at 42°C, ER stress: 2 hours with 1μM Thapsigargin, Proteasome inhibition: 2 hours with 1μM MG132. As a control, 2 hours with 1μl DMSO per 1ml medium was used.

### GFP-Tagged proteins

Plasmids expressing GFP-tagged proteins were prepared using the pcDNA3.1 backbone plasmid. RPLP0, RPS2, RPL28, RPL10A, and BTF3 proteins were tagged with an N-terminal GFP tag and cloned into pcDNA3.1 using Gibson cloning. HSPA14 protein was tagged with a C-terminal GFP tag, or with a C-terminal FLAG tag using the pcDNA3.1 backbone plasmid, and with a N-terminal GFP tag using a pT-Rex™-DEST30 plasmid, using gateway cloning.

### GFP intensity measurement and Fluorescent microscopy

To measure the expression level of GFP-tagged RPs, GFP fluorescence intensity of cells was measured in live cells, after replacing media with PBS, using Tecan Infinity M200PRO reader. Non-transfected cells were used as background.

Fluorescent microscopy imaging of HEK293T cells transfected with GFP-tagged ribosomal proteins was performed with the following exposure times for GFP channel: RPL10A - 400ms, RPL28 - 1000ms, RPLP0 - 200ms, RPS2 - 200ms.

### Sucrose gradient preparation

Sucrose solutions were prepared according to the following recipe: 25mM HEPES pH 6.9, 100mM KCl, 5mM MgCl2, and 10% or 50% sucrose. Protease inhibitors (Roche tablets cat #11836170001), 100μg/ml CHX (cycloheximide), and 1mM DTT (dithiothreitol) were added immediately before use.

Sucrose gradients of 10%-to-50% sucrose were prepared using BioComp gradient master, according to the manufacturer instruction.

### Cell harvesting for fluorescent polysome profiling

Lysis buffer was prepared as follows: 25mM HEPES pH 6.9, 1% Triton X-100, 100mM KCl and 5mM MgCl2. Protease inhibitors (Roche tablets, cat #11836170001), 100μg/ml CHX, 1mM DTT, and RNase inhibitors (SUPERase-In, ThermoFisher, AM2694) were added immediately before use. Cells were incubated with 100μg/ml CHX for 3 minutes at 37 °C, then put on ice, washed once with ice cold PBS with 100μg/ml CHX and collected by scraping in ice cold PBS with 100μg/ml CHX. After pelleting at low speed (300g for 7 minutes at 4°C), cells were resuspended in ice cold lysis buffer. Cellular membranes were disrupted by passing through a 25G syringe needle 5 times. Then, the lysates were centrifuged for 10min at 15000 RPM at 4 °C, and the supernatant was transferred to new tubes. OD units (260nm) were measured using NanoDrop 2000C spectrophotometer.

### Fluorescent Polysome Profiling

Sample volumes containing equal number of 260nm OD units (corresponding to the amount of RNA per sample) were loaded onto the sucrose gradients, and centrifuged using an ultracentrifuge for 2 hours at 35,000 RPM at 4°C. UV and GFP polysome profiles were obtained using the Triax™ flowcell (BioComp Instruments). Triax™ flowcell software version 1.45A was used with the following parameters:

| **Parameter** | **Value** |
| --- | --- |
| Scanning mode | UV OD WITH SINGLE FLUORESCENCE SCAN |
| Channel A (LED1) Wavelength | 260 nm |
| Channel B (LED2) Fluor | EGFP |
| Excitation (LED2) Wavelength | 470 nm |
| Excitation filter A | 470/40 nm |
| Emission filter A | 525/50 nm |
| Sensitivity | 0 |
| Integration time | 350 ms |
| LED1 (260 nm) On Time | 18 ms |
| PD2 (470 nm) On Time | 350 ms |

A sample of equal amount of 260 nm OD units from non-transfected HEK293T cell lysate was ran together with the other experimental samples. This OD unit matched negative control GFP fluorescence profile was used as background, which was subtracted from every other GFP fluorescence profile with the same 260nm OD unit loaded. We note that subtraction of an equally loaded 260 nm OD unit fluorescent polysome profile of nontransfected cell lysate is essential to obtain a reliable quantification of the polysome-association of fluorescently tagged proteins.

All fluorescent polysome profile figures were prepared using MATLAB scripts.

For display purposes polysome curves were smoothed using a median filter, with a window of 10 for UV curves and a window of 15 for GFP curves.

### Quantification of ribosomal association changes from Fluorescent Polysome Profiles

In order to calculate the changes in the polysome-association of a fluorescently-tagged protein, the following method was employed. First, for all fluorescence curves, the background (non-transfected sample) fluorescence profile was subtracted. Then, for each sample (Control and stress), the UV curve and the fluorescence curve were each divided into four regions: Monosome (single ribosome per mRNA), Low polysomes (2 to 5 ribosomes per mRNA), Medium polysomes (6 to 8 ribosomes per mRNA) and High polysomes (9 or more ribosomes per mRNA). Area under the curves were calculated for each region, and the area of each fluorescence region (representing interaction) was divided by the corresponding UV area (representing polysome amounts), resulting in normalized ribosome association per region along the gradient. Finally, to estimate the fold change in polysome association in response to stress, normalized ribosome association values in stress samples were divided by their corresponding normalized ribosome association values in the Control sample for each region.

### Polysomal fractionation followed by WB

Cells transfected with HSPA14-FLAG, either untreated or subjected to HS, were harvested and polysome profiling was performed as above. In parallel, polysomal fractions were collected using a Gilson FC203B fraction collector, with 24 equal volume (450μl) fractions per condition. Following polysome fractionation, the fractions were combined into five pools, the Unbound fraction (fractions 1-3), ribosome Subunits (Sub, 4-6), Monosome (Mono, 7-8), Medium polysomes (Med, 9-14) and Heavy polysomes (Heavy, 15-22). The Sub, Mono, Med and Heavy pools were each diluted to a final volume of 12.5ml with a dilution buffer (20mM HEPES pH 6.9, 0.5% NP-40, 0.5% Sodium Deoxycholate, 100mM KCl and 5mM MgCl2. 100μg/ml CHX was added immediately before use). Then, the Sub, Mono, Med and Heavy pools were layered on top of 12.5ml of sucrose cushion solution (25mM HEPES pH 6.9, 100mM KCl and 5mM MgCl2, and 17.5% sucrose. 100μg/ml CHX was added immediately before use) and centrifuged for 17 hours at 44,000RPM in a type 70 Ti rotor, at 4°C (an equivalent of 100,000G for 24 hours, as in (Rivera, Maguire et al., 2015)). Subsequently, the remaining ribosome pellets were resuspended in 40ul of protein sample buffer. The Unbound pool was diluted 1:1 with protein sample buffer, and all samples were boiled at 100°C for 5 minutes. Finally, equal volumes of each fraction pool, corresponding to 0.5% of the Unbound fraction and 37.5% of each of the other fractions, were loaded on a 10% SDS-PAGE gel and immunoblotted with anti-FLAG (Sigma-Aldrich, F1804) and anti-RPL10A (Abcam, ab174318) antibodies.

### Co-immunoprecipitation

Co-immunoprecipitation (co-IP) buffer was prepared as follows: 50mM HEPES pH 7.9, 0.5% Triton X-100, 5% glycerol, 150mM NaCl and 5mM MgCl2. Protease inhibitors (Roche tablets, cat# 11836170001), 100μg/ml CHX, 1mM DTT and RNase inhibitors (SUPERase-In, ThermoFisher, AM2694) were added immediately before use. Wash buffer was prepared as follows: 50mM HEPES pH 7.9, 1% Triton X-100, 5% glycerol, 150mM NaCl and 5mM MgCl2. Cells were harvested as for fluorescent polysome profiling described above, using co-IP buffer for lysis. For RNAse treatment, 1μl RNase If (NEB, M0243) per one 260 nm OD unit was added, and samples were rotated for 55 min at room temperature. Then 2μl of SUPERase-In (ThermoFisher, AM2694) RNase inhibitor were added to all samples. Samples were immunoprecipitated using 20μl of EZview Red ANTIFLAG M2 Affinity Gel beads (Sigma-Aldrich, F2426), which were added to a sample volume containing equal number of 260 nm OD units. Additional co-IP buffer was added to reach a final volume of 1ml per sample. Samples with beads were slowly rotated overnight at 4 °C. Beads were then washed 7 times with 1 ml of wash buffer. Following the last wash, the beads were resuspended in equal volumes of wash buffer, protein sample buffer was added, and the samples were boiled at 100 °C for 5 minutes. Equal volumes were then loaded on an 8% NuPAGE gel, and immunoblotted using anti-FLAG (Sigma-Aldrich, F1804) and anti-GFP (MBL, 598) antibodies.

### Western Blot quantification

Quantification of the Western Blot bands was performed using Fiji software (Schindelin, Arganda-Carreras et al., 2012). To calculate the fold change in co-immunoprecipitation of GFP-labeled HSPA14 with HSPA14-FLAG, ratios of anti-GFP and anti-FLAG bands were calculated for each sample. Then, a ratio of the Heat Stress to Control ratios was calculated to calculate the fold change as a result of stress.

## Supplementary data

Supplementary Information file contains Supplementary Figures S1-S4.

## Acknowledgement

We thank Jeeda Bisharat for her help throughout the project, and Oded Lewinson for helpful discussions. We thank Herman Wolosker, Flonia Levy-Adam and Kinneret Rozales for critical reading of the manuscript.

## Authors Contributions

R.S. conceived the study. R.S. and A.M. designed the experiments. D.C. developed the Triax flowcell instrument. A.M. performed all the experiments. R.S. wrote the manuscript with input from A.M.

