## Supplementary Information for "Fluorescent polysome profiling reveals stress-mediated regulation of HSPA14-ribosome interactions" (from another version of this preprint: version 1, 2019-12-01)

[2] BioComp Instruments. Canada

### Supplementary Figures

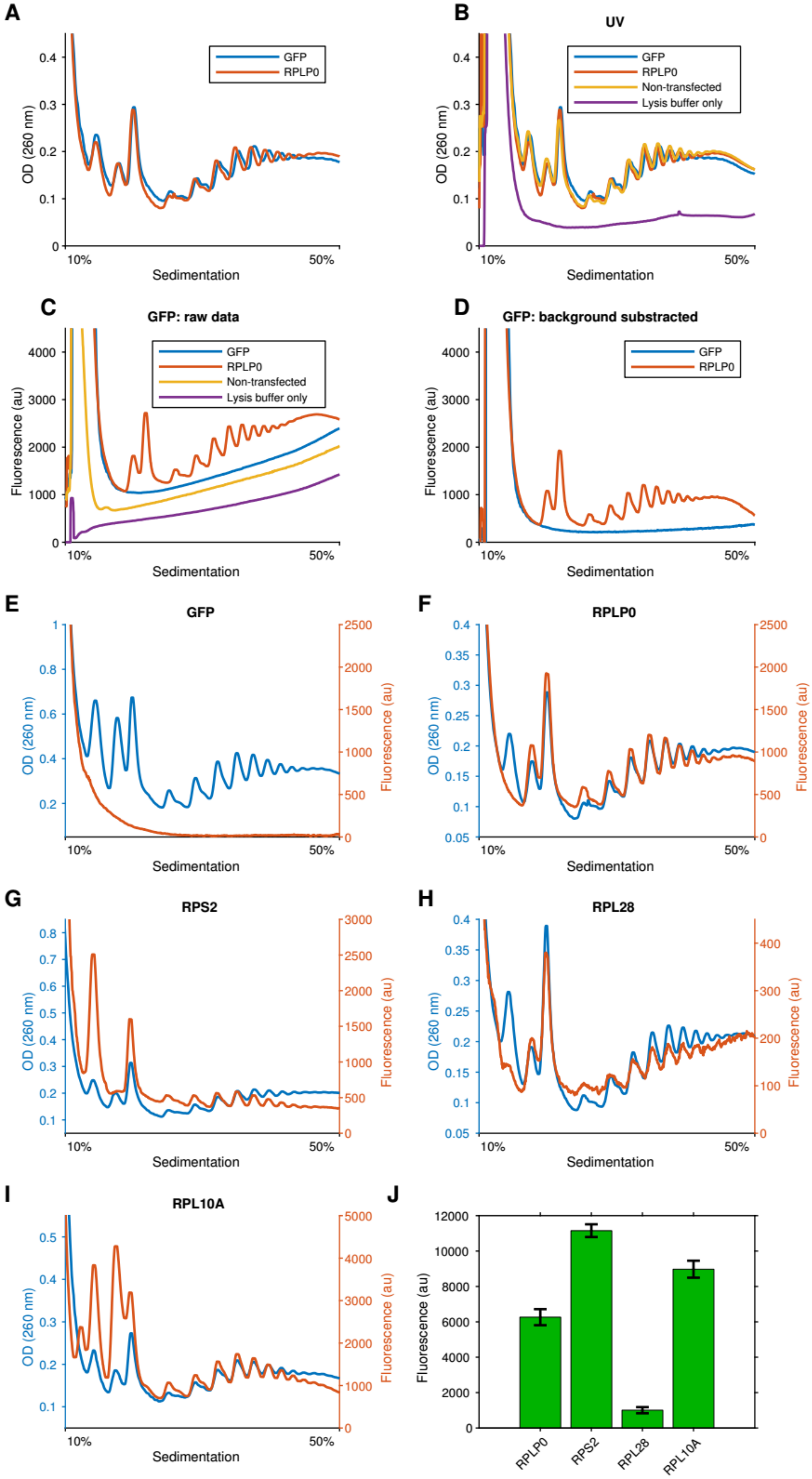
A
GFP
RPLP0
OD (260 nm)
Sedimentation
10%
50%
B
UV
GFP
RPLP0
Non-transfected
Lysis buffer only
C
GFP: raw data
Fluorescence (au)
D
GFP: background substracted
E
GFP
F
RPLP0
G
RPS2
H
RPL28
I
RPL10A
J
RPLP0
RPS2
RPL28
RPL10A

**Figure S1: Fluorescent polysome profiling reliably detects GFP-tagged ribosome-incorporated ribosomal proteins.**

(A) UV polysome profiles of GFP-only or GFP-tagged RPLP0 transfected HEK293T cells shows no significant difference in the polysome levels and distribution. (B) UV polysome profiles of non-transfected HEK293T cells, GFP-only or GFP-tagged RPLP0 transfected HEK293T cells shows no significant difference in poylsome levels and distribution, indicating that the transfection and ectopic expression of GFP or GFP tagged RPLP0 did not affect translation. Lysis buffer only UV reads are also shown. (C) Raw data of fluorescent polysome profiles of non-transfected HEK293T cell lysates, or lysis buffer only shows the level of fluorescence background along the gradient. GFP-only or GFP-tagged RPLP0 transfected cell lysate raw fluorescence profiles from the same run are shown in comparison. (D) GFP fluorescence polysome profiles of GFP-only or GFP-tagged RPLP0 transfected cells, after subtraction of the fluorescence background profile of non-transfected HEK293T cell lysates. (E-I) Independent biological replicates for Fig. 1B-E, fluorescent polysome profiling of GFP only (E), GFP-tagged RPLP0 (F), RPS2 (G), RPL28 (H) and RPL10A (I). All fluorescent profiles were background subtracted using non-transfected HEK293T lysates from the same run with equal OD units. (J) GFP fluorescence intensity in cells expressing GFP-tagged RPLP0, RPS2, RPL28 or RPL10A was measured using a fluorescent reader (Tecan M200, see Methods) to illustrate different expression levels of GFP-tagged RPs.

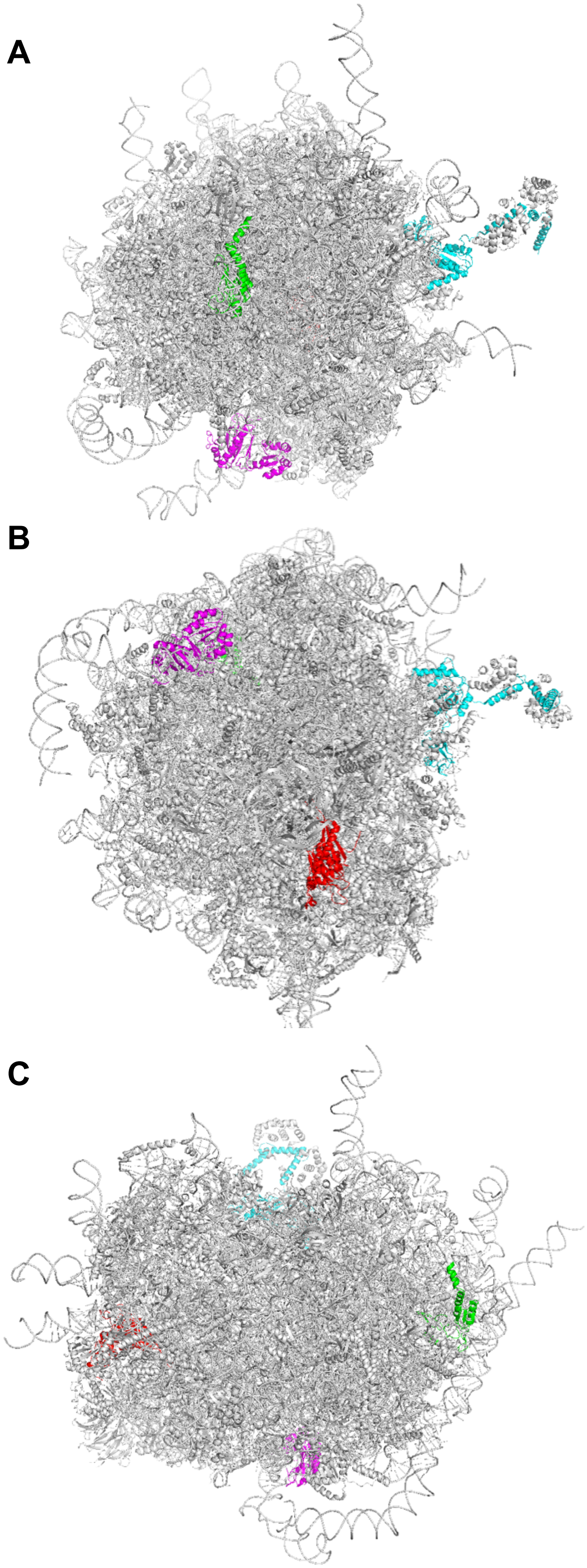
A
B
C

**Figure S2: Chosen Ribosomal Proteins (RPs) are surface accessible.**

Locations of RPS2 (red), RPL10A (pink), RPLP0 (cyan) and RPL28 (green) on the ribosome. Ribosome structure was obtained from the PDB database (accession number 4v6x, Anger et al. [1]), and structures drawn using PyMol.

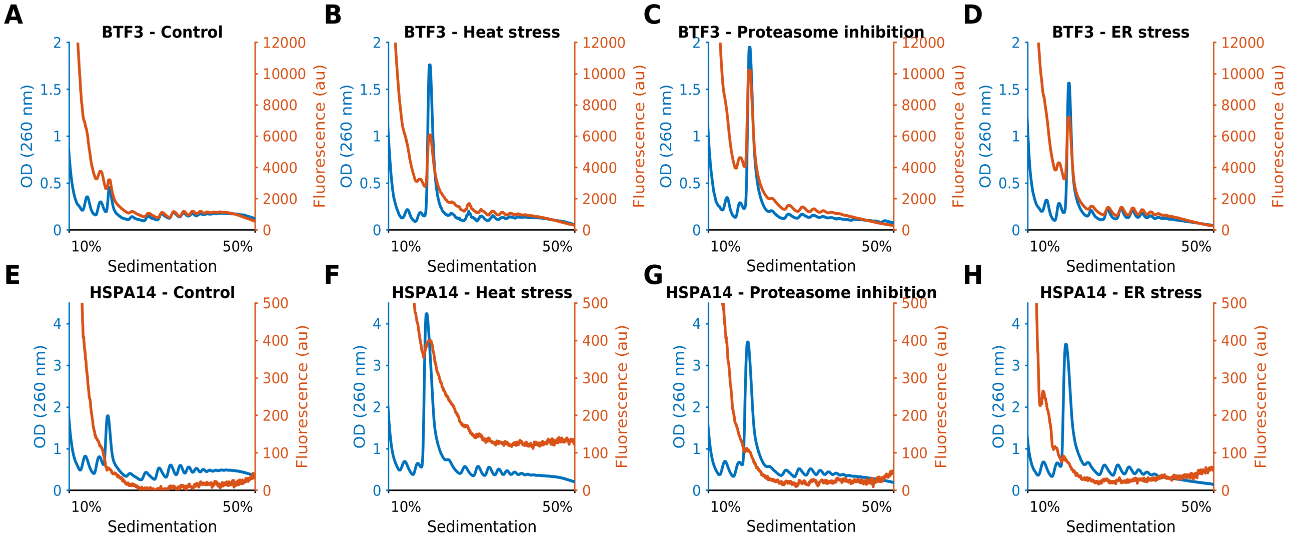


**Figure S3: Fluorescent polysome profiling of ribosome-associated chaperones in response to stress.**

(A-D) GFP-tagged BTF3 protein ribosome association before and after stress. Independent biological replicate of Fig. 2A-D. (E-H) GFP-tagged HSPA14 polysome association before and after stress. Independent biological replicate of Fig. 3A-D.

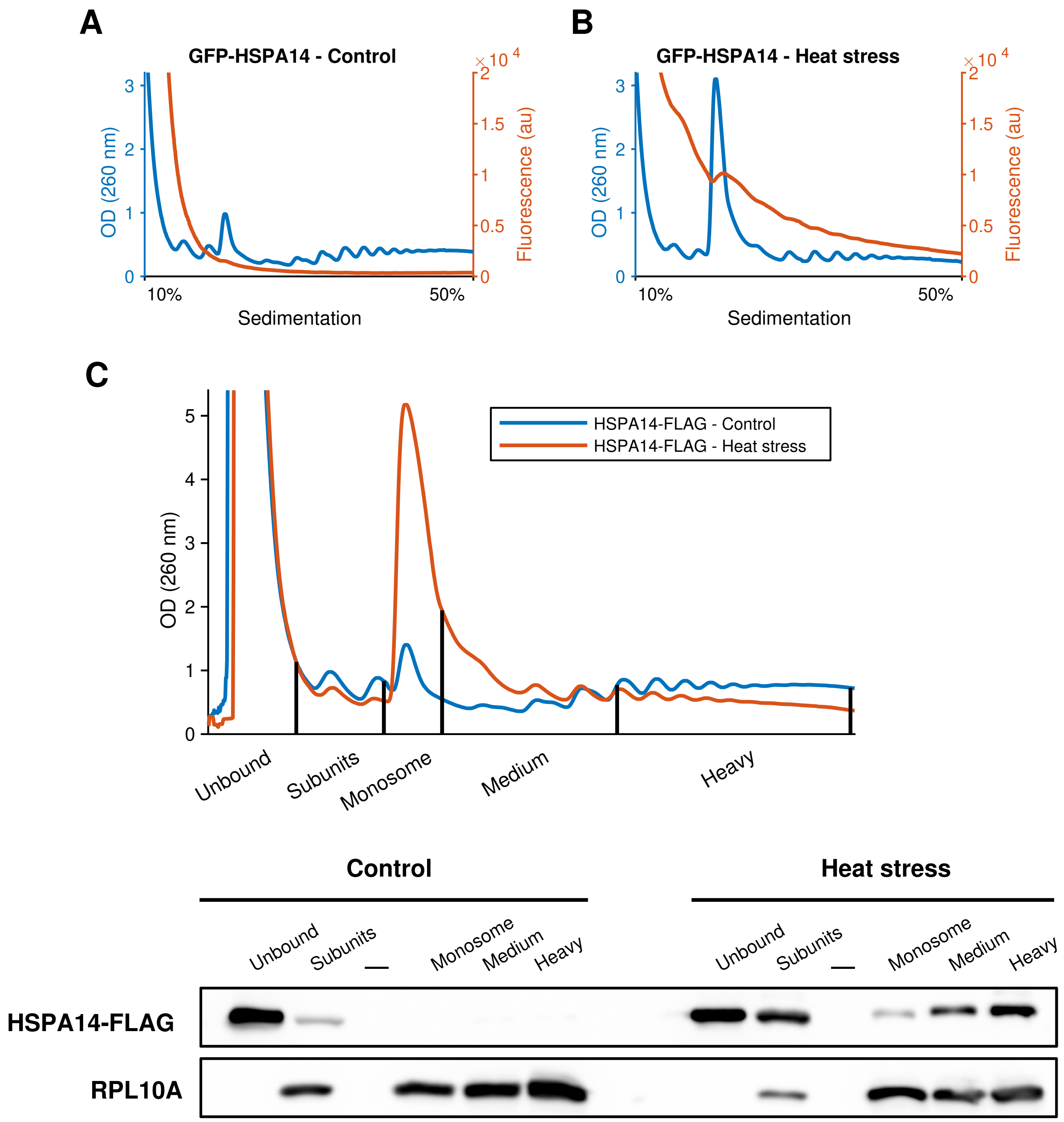


**Figure S4: HSPA14 ribosome association in Heat stress is tag-independent.**

(A-B) N-terminal GFP-tagged HSPA14 shows a marked increase in polysome interactions following heat stress (B) compared to control conditions (A) using fluorescent polysome profiling, similarly to the C-terminal GFP-tagged HSPA14 (Fig. 3). An independent biological replicate of Fig. 4A,B. (C) Polysome profiling fractionation followed by protein extraction and WB of cells transfected with HSPA14-FLAG. Independent biological replicate of Fig. 4C.

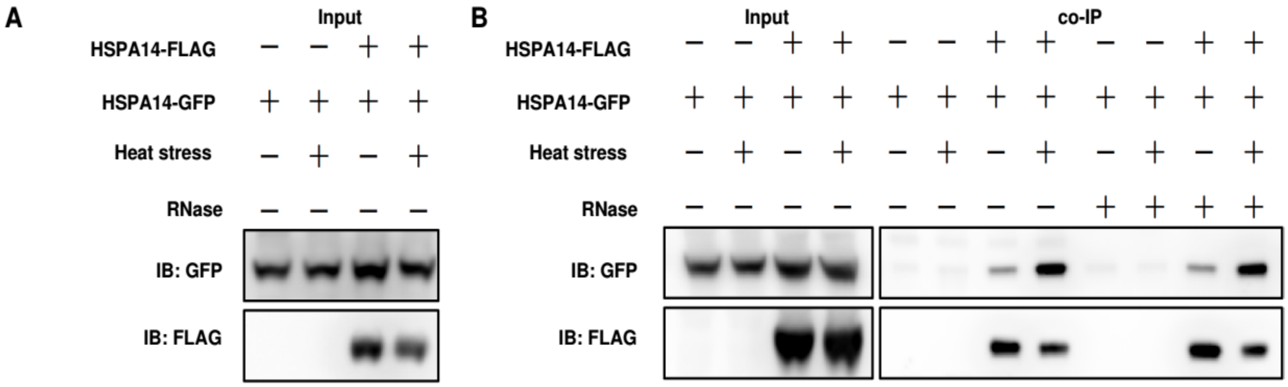


**Figure S5: HSPA14-HSPA14 interaction increases in response to heat shock.**

**(**A) Input immunoblot of the co-immunoprecipitation (co-IP) from Fig. 4D. (B) Co-IP reveals an increased HSPA14-HSPA14 interaction following heat stress, an independent biological replicate for Fig. 4D.
